# 16SpeB: Towards defining bacterial species boundaries by intra-species gene sequence identity

**DOI:** 10.1101/060657

**Authors:** Adam Chun-Nin Wong, Patrick Ng, Angela E. Douglas

## Abstract

**Summary:** 16SpeB (<u>16</u>S rRNA-based <u>Spe</u>cies <u>B</u>oundary) is a package of Perl programs that evaluates total sequence variation of a bacterial species at the levels of the whole 16S rRNA sequences or single hypervariable (V) regions, using publicly-available sequences. The 16SpeB pipelines filter sequences from duplicated strains and of low quality, extracts a V region of interest using general primer sequences, and calculates sequence percentage identity (%ID) through all possible pairwise alignments.

**Results:** The minimum %ID of 16S rRNA gene sequences for 15 clinically-important bacterial species, as determined by 16SpeB, ranged from 82.6% to 99.8%. The relationship between minimum %ID of V2/V6 regions and full-gene sequences varied among species, indicating that %ID species limits should be resolved independently for each region of the 16S rRNA gene and bacterial species.

**Availability:** 16SpeB and user manual are freely available for download from: https://github.com/pnpnpn/16SpeB. A video tutorial is available at: https://youtu.be/Vd6YmMhyBiA

**Supplementary information:** Supplementary data are available at Bioinformatics online.

## 1 INTRODUCTION

Fueled by recent advance in next-generation sequencing (NGS), nucleic-acid-based identification of microbes from clinical and environmental samples is an emerging area of scientific interests (Kress, et al., 2015; Shokralla, et al., 2012; van Dijk, et al., 2014; Wilson, et al., 2014). For bacteria, gene markers such as the 16S ribosomal RNA (rRNA) gene are commonly used to profile communities that encompass both cultured and uncultured species. An enduring challenge is to assign taxonomy to these marker gene sequences, especially, to assess the confidence a particular sequence read fits into its designated taxonomic rank based on percentage identity (%ID); and be able to discriminate rare or novel taxa from taxa likely arisen from sequencing errors.

Over time, scientists have attempted to find a unifying threshold to define bacterial species boundary from their gene sequences. For example, a 97% sequence identity (%ID) of the full length 16S rRNA gene has been put forward as the cut-off value to define species (Drancourt, et al., 2004; Drancourt and Raoult, 2005; Ueda, et al., 1999), but the criterion has been vigorously challenged (Clarridge, 2004; Janda and Abbott, 2007; Petti, 2007; Rossi-Tamisier, et al., 2015). Compounding the uncertainty about using a fixed %ID threshold for species identification, it is becoming a common trend to sequence shorter but varied reads (<400 bp) of single hypervariable (V) regions, such as the V2 or V6 of the 16S rRNA gene (Bowen, et al., 2011; De Filippo, et al., 2010; Guss, et al., 2011; Kirchman, et al., 2010; Ravussin, et al., 2011; Werner, et al., 2012; Wu, et al., 2011).

To address some of the caveats associated with 16S rRNA gene profiling, especially to facilitate more confident taxonomy assignment, we proposed that the 16S rRNA %ID variation from known sequences shall be determined and used to guide the boundary of bacteria species-to-species. We thus develop 16SpeB (<u>16</u>S rRNA-based <u>Spe</u>cies <u>B</u>oundary). 16SpeB is an analytical tool designed to identify the range of 16S %ID encompassed by individual bacterial species based on known 16S rRNA gene sequence variation. Our goal is to promote accurate taxonomic identification of bacteria in both (near)-full 16S sequences and short reads obtained by 454, Illumina or other next-generation sequencing platforms.

## 2 USAGE

16S rRNA sequences from three 16S rRNA databases can be downloaded from *Greengenes* (DeSantis, et al., 2006) *Ribosomal Database Project* (Cole, et al., 2007) and *Silva* (Pruesse, et al., 2007). 16SpeB allows users to trim the (near-)full 16S rRNA sequences to their preferred length. It can also extract the sequences of the V2 and V6 regions, which are widely used in 454 sequencing studies, by reference to the general primer sets 27F-338R and 784F-1061R, respectively. Sequences that fail to satisfy the two following conditions are removed: (1) <2 bp mismatches with the general 16S primers (i.e. conserved regions of the 16S gene), and (2) relative coordinates of matched primers are within +/− 50 bp from the relative coordinates of the literature. The V2 region is trimmed to 270 bp upstream of the 338R primer. 16SpeB conducts all possible pairwise sequence comparisons by aligning all pairwise sequences using Needleman-Wunsch alignment algorithm with match/mismatch score of 1/−2 and affine gap penalty open/extension of −5/−2. The minimum and 95% quantile %ID are computed for each species, providing a measure of the total known sequence variation that defines the species.

## 3 APPLICATION OF 16SpeB

16SpeB was initially developed to identify species limits of *Acetobacter* and *Lactobacillus* in a pyroseqeuncing analysis of the gut microbiota of *Drosophila melanogaster* (Wong, et al., 2011). Here we extend the application of 16SpeB to determine the %ID of (near-)full 16S rRNA genes that defines the species boundary of 15 clinically-important bacterial species (listed in Supplementary Data Set 1); and to determine the %ID of the V2 and V6 regions widely used in pyrosequencing studies that correlate with this species boundary. The 15 bacterial species were selected on the criteria that a broad range of publicly-available sequences (3 to 454) and phylogenetic diversity (including representatives of Actinobacteria, Bacteroidetes, Chlamydiae, Firmicutes and Proteobacteria) were represented. In total, 1,296 sequences were analyzed. The minimum %ID of (near-) full 16S sequences varied from 99.8% (*Neisseria gonorrhoeae*) to 82.6% (*Staphylococcus aureus*) (Table 1). Just two (13%) of the 15 species had minimum %ID close to predicted 97% threshold for species boundary (*Neisseria meningitidis* 97.0%, and *Listeria monocytogenes* 97.1%); and 11 (73%) species deviated from 97% by more than one percentage point. Values of the 95% quantile are provided in Table 1 and may prove to be more useful than minimum %ID for some species, e.g. *Staphylococcus aureus,* where the minimum %ID is suspected to be artefactually low (possibly through mis-identification).

**TABLE 1.** The minimum and 95% quantile %ID of the (near-)full 16S rRNA gene, and the V2 and V6 regions of the 15 clinically-important bacteria.

| Species | Number of sequences<br>(pairs) | Minimum %ID |  |  | 95% quantile %ID |  |  |
| --- | --- | --- | --- | --- | --- | --- | --- |
|  |  | (near)-full<br>16S | V2 | V6 | (near)-full<br>16S | V2 | V6 |
| *Bacteroides fragilis* | 345 (59340) | 0.928 | 0.899 | 0.928 | 0.978 | 0.959 | 0.973 |
| *Clostridium bifermentans* | 58 (1653) | 0.926 | 0.928 | 0.787 | 0.967 | 0.967 | 0.893 |
| *Chlamydia trachomati* | 15 (105) | 0.950 | 0.921 | 0.954 | 0.958 | 0.942 | 0.959 |
| *Corynebacterium diphtheriae* | 10 (45) | 0.942 | 0.920 | 0.912 | 0.946 | 0.934 | 0.918 |
| *Haemophilus influenzae* | 92 (4186) | 0.901 | 0.831 | 0.891 | 0.951 | 0.925 | 0.907 |
| *Helicobacter pylori* | 59 (1711) | 0.949 | 0.895 | 0.939 | 0.977 | 0.960 | 0.961 |
| *Listeria monocytogenes* | 26 (325) | 0.971 | 0.939 | 0.966 | 0.974 | 0.953 | 0.969 |
| *Mycobacterium leprae* | 4 (6) | 0.984 | 0.967 | 0.992 | 0.984 | 0.967 | 0.992 |
| *Mycobacterium tuberculosis* | 10 (45) | 0.984 | 0.982 | 0.988 | 0.989 | 0.985 | 0.992 |
| *Mycoplasma hominis* | 6 (15) | 0.899 | 0.949 | 0.681 | 0.899 | 0.949 | 0.681 |
| *Neisseria gonorrhoeae* | 3 (3) | 0.998 | 1.000 | 1.000 | 0.999 | 1.000 | 1.000 |
| *Neisseria meningitidis* | 133 (8778) | 0.970 | 0.927 | 0.962 | 0.990 | 0.978 | 0.981 |
| *Staphylococcus aureus* | 454 (102831) | 0.826 | 0.604 | 0.843 | 0.980 | 0.981 | 0.973 |
| *Streptococcus pneumoniae* | 47 (1081) | 0.980 | 0.938 | 0.977 | 0.986 | 0.963 | 0.985 |
| *Yersinia pestis* | 34 (561) | 0.979 | 0.960 | 0.966 | 0.986 | 0.967 | 0.977 |

As anticipated, the minimum %ID of both the V2 and V6 regions varied positively with %minimum ID of the (near-) full sequence of the 16S genes (Supplementary Figure 1). The relationships were not, however, tight indicating that the rates of sequence evolution of individual V regions are not closely correlated to each other or to other regions of the 16S gene. The implication is that, just as the 97% threshold is not a reliable index of the taxonomic species limit, so there is no simple linear relationship linking the minimum %ID of the V2 or V6 sequences to the (near-) full 16S sequence across multiple bacterial species.

We conclude that the %ID species limits should be resolved independently for each region of the 16S rRNA gene and each bacterial species. Therefore, 16SpeB can serve as an important tool that facilitates accurate taxonomic identification and proper interpretation of 16S rRNA gene pyrosequencing data.

## Acknowledgements

The project described was supported by Grant Number R01GM095372 from the National Institute of General Medical Sciences (NIH), and by Sarkaria Institute of Insect Physiology and Toxicology. The content is solely the responsibility of the authors and does not necessarily represent the official views of the National Institute of General Medical Sciences or the National Institutes of Health.

SUPPLEMENTARY DATA SET 1. List of 16S rRNA 118 sequences used in the study

SUPPLEMENTARY FIGURE 1. Relationship between a) minimum and b) 95% quantile %ID of V2/V6 region and (near)-full 16S rRNA gene sequence across the 15 bacterial species used in this study. (V2 region: black, solid squares; V6 region: grey, open circles).

